# Progerin-Induced Nuclear Envelope Remodeling is Shaped by Cell Division and NUP153

**DOI:** 10.64898/2026.01.23.701344

**Authors:** Ayse Mubine Turkmen, Jing Wang, Adam Frost, Katharine S. Ullman

## Abstract

The nuclear envelope (NE) undergoes dynamic remodeling during both physiological and pathological processes. Nuclei in cells from patients with accelerated aging diseases and from elder individuals are often lobular and convoluted. Despite extensive study, the steps of phenotype acquisition and the co-factors required are not yet fully understood. Here, we focused on progerin, a mutant form of lamin A that causes Hutchinson Gilford Progeria Syndrome (HGPS). Using an inducible cell-based system, we characterized two distinct stages of NE remodeling. Correlative light and electron microscopy of interphase-arrested cells showed that prior to cell division, progerin primarily affects the inner nuclear membrane (INM), inducing focal expansion, invagination, and the formation of multi-membranous structures, while the outer nuclear membrane remains largely unaffected. These focal regions of progerin accumulation are enriched for specific INM proteins and the nucleoporin NUP153 but largely exclude nuclear pore complexes. Live and fixed image analysis demonstrated that, upon cell division and NE reassembly, progerin-expressing cells develop pronounced nuclear lobulations characteristic of HGPS. Appreciation of this stepwise development of phenotype lends insight into cellular manifestations of aging in mitotic versus post-mitotic cell types and provides a system in which to study factors that contribute to these distinct stages in the disruption of nuclear morphology. Depletion of NUP153 reduced NE foci formation in interphase-arrested cells expressing GFP-progerin, suggesting NUP153 promotes or stabilizes INM invagination. Aberrant nuclear architecture is just one cellular feature that changes during normal and accelerated aging but its etiology provides a critical framework for understanding accompanying consequences on chromatin packaging, DNA damage, and the ER stress response.

## INTRODUCTION

In eukaryotic cells, genomic DNA is enclosed by the nuclear envelope (NE) a specialized membrane system comprised of an outer nuclear membrane (ONM) and inner nuclear membrane (INM). The INM houses a distinct repertoire of transmembrane proteins critical to the organization of underlying nuclear lamina and chromatin while the ONM is continuous with the endoplasmic reticulum and shares many of its components. Certain proteins, such as nesprins, are enriched at the ONM via luminal domain interactions with INM proteins (Crisp et al., 2006; Padmakumar et al., 2005). Nuclear pore complexes (NPCs) are macromolecular structures where the ONM and INM join at the site of proteinaceous channels that, together with soluble factors, mediate traffic between cytoplasm and nucleoplasm.

Despite the static appearance of a canonical ovoid nucleus at interphase, NE architecture is highly dynamic (Chu et al., 2017; Keuenhof et al., 2023; McPhee et al., 2024; Turkmen et al., 2023). Overt changes in NE organization occur during mitosis where the NE disassembles and reforms in mammalian cells (Beaudouin et al., 2002; Ulbert et al., 2006). Beyond mitotic dynamics, NE architecture exhibits other forms of structural remodeling. The NE can fold inward linearly to form shallow clefts or develop tubular invaginations of the INM or the INM and ONM together, creating narrow membrane-enclosed channels within the nucleus termed type I and type II nucleoplasmic reticulum (NR) respectively (Malhas et al., 2011; Stiekema et al., 2022). At times, these involutions traverse from one side of the nucleus to the other, forming a tubule. Such fluctuations in NE structure play crucial roles in diverse physiological processes such as gene expression (Biedzinski et al., 2020; Feurle et al., 2021) and DNA repair (Legartová et al., 2014; Shokrollahi et al., 2024) but can also be aberrantly heightened under pathological conditions (Fischer, 2020; Frost et al., 2016; Janin et al., 2017; Todosenko et al., 2025).

Numerous disease-associated alleles of NE and associated proteins result in nuclear dysmorphia, underscoring the importance of this environment for cell function (Cabanillas et al., 2011; Chen et al., 2023; Janin et al., 2017). Lamin A (*LMNA*), a component of the nuclear lamina, has multiple disease-associated alleles, including the mutation that results in the rapid aging disease, Hutchinson-Gilford Progeria Syndrome (HGPS) (Shin and Worman, 2022). In HGPS, a point mutation activates a cryptic splice site in the *LMNA* gene, producing a lamin A mutant that lacks the proteolytic cleavage site needed to remove its farnesylated C-terminus during maturation. This lamin A isoform, termed progerin (Fig. S1A), is also produced in tissues of aged individuals through spontaneous cryptic splice site usage, suggesting a role in physiological aging (Goulbourne et al., 2011; McClintock et al., 2007; Olive et al., 2010; Revêchon et al., 2025; Rodriguez et al., 2009). While multiple factors contribute to aging, progressive deterioration of nuclear architecture is a hallmark feature (Pathak et al., 2021). In patient-derived HGPS cells, prominent features are pronounced nuclear folds and lobulations (Scaffidi and Misteli, 2006) that have been noted to worsen with passage number (Goldman et al., 2004; Röhrl et al., 2021).

To study how progerin expression results in NE remodeling, we focused on early steps of phenotype development following temporally restricted progerin production. This analysis revealed distinct stages: first a tubular-lamellar expansion of the INM forms while overall NE shape is maintained and then, following cell division and nuclear reformation, an expanded and lobulated NE arises. The appreciation of different phases in the progression of NE deformation allowed us to isolate early events in phenotype acquisition for further study and create a testing ground for contributing factors.

## RESULTS AND DISCUSSION

### Development of the characteristic HGPS nuclear phenotype requires mitosis

To investigate the events of NE remodeling in response to progerin expression, we employed a doxycycline-inducible GFP-progerin expression system in U2OS cells and performed time-lapse confocal microscopy. GFP-progerin localized to the nuclear rim approximately 5 hours post-doxycycline induction, demarcating a regular ovoid shape, and over time discrete GFP-positive (GFP⁺) foci emerged (Fig. 1A, white arrows). These foci were clearl y associated with the NE (Fig. S1B). As expression continued, the number and intensity of GFP⁺ foci increased and nuclear shape was still regular. Notably, by 11 hours post-induction, only cells that had undergone mitosis and NE reassembly exhibited irregular nuclear morphology with pronounced folds and lobulations (Fig. 1A, yellow arrowheads). This suggests that progerin initially targets to the nuclear rim while also accumulating at discrete nuclear foci and then, following NE reassembly post mitosis, becomes distributed primarily at the nuclear periphery as the nucleus concomitantly develops a highly lobulated and folded appearance characteristic of HGPS.

**Figure 1.**
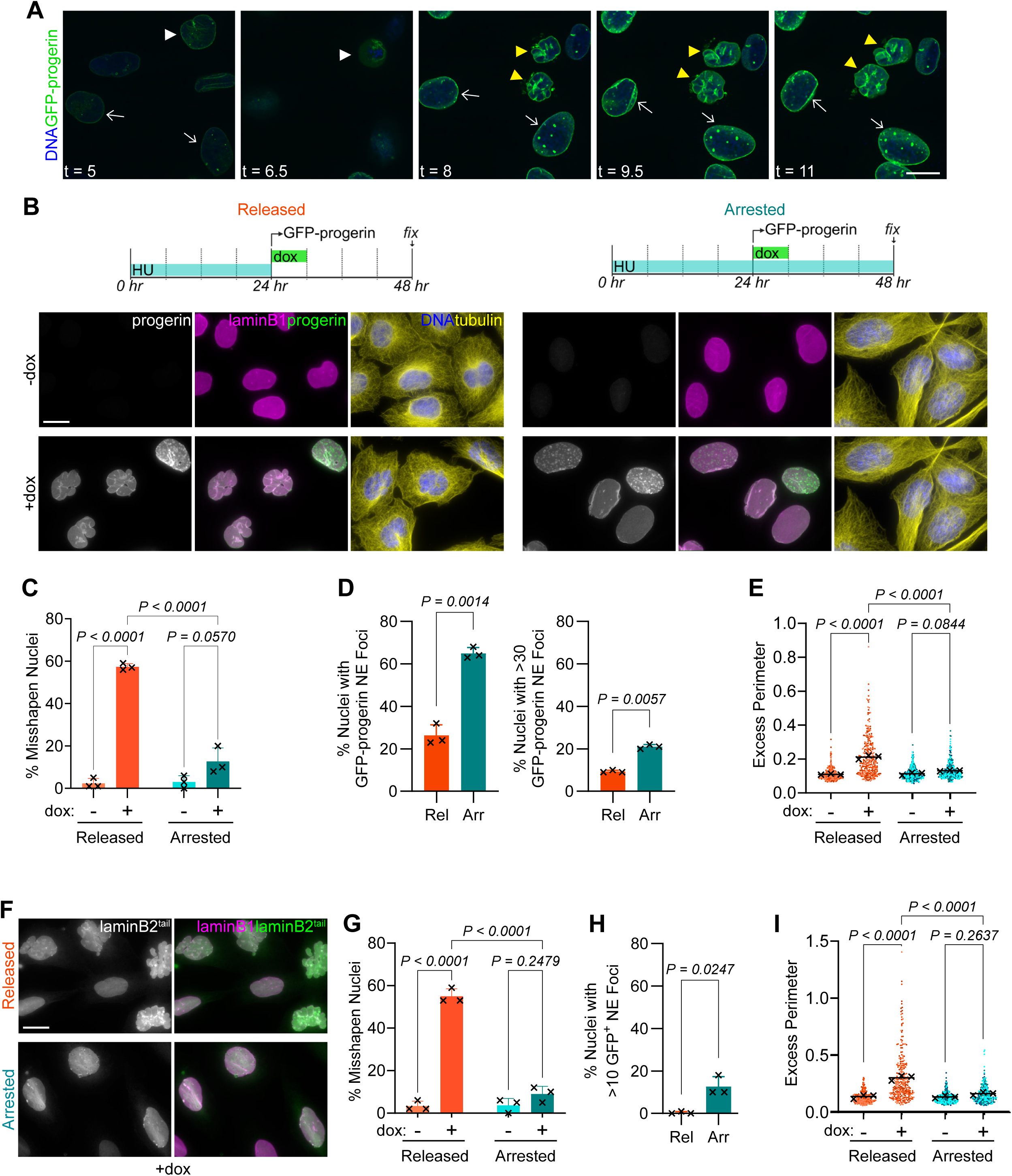
The lobulated nuclear phenotype associated with HGPS is dependent on cell division, with arrested cells displaying a distinct nuclear phenotype. 1A. Time-lapse imaging of U2OS cells with dox-inducible GFP-progerin expression. Cells were incubated with 200ng/ml dox at t = 0. Images were acquired every 30 minutes for 7 hours using a spinning disc confocal microscope. DNA was visualized using NucBlue™. White arrows: nuclei of cells that have not divided; arrowheads: pre-division nucleus (white) and nuclei post mitosis (yellow). Scale bars in Figure 1 are 20 μm 1B. Immunofluorescence widefield microscopy of U2OS cells with dox-inducible GFP-progerin expression in “Released” and “Arrested” experimental conditions. Cells were stained for lamin B1, tubulin, and GFP (progerin) and NucBlue to detect DNA. 1C. Percent of misshapen nuclei from experiments described in 1B. Nuclei with convexity less than 0.945 were classified as misshapen. Bars represent the mean of three experiments; error bars represent standard deviation; each X represents a technical replicate. n=3, N=100. Data was analyzed using a two-way ANOVA with post-hoc Tukey’s test. 1D. Percent of nuclei from dox-treated cells in the experiments described in 1B with any GFP^+^ foci (left) or >30 GFP^+^ foci (right). Bars represent the mean of three experiments; error bars represent standard deviation. Each X represents the mean of a technical replicate. n=3, N=100. Data was analyzed using paired two-tailed t-tests. Rel = released; Arr = arrested 1E. Excess nuclear perimeter from experiments described in 1B. Dots represent single nuclei, with different shades representing replicates; each X represents the mean of a technical replicate. n=3, N=100. Data was analyzed using a two-way ANOVA with with post-hoc Tukey’s test. 1F. Widefield microscopy of U2OS cells with dox-inducible GFP-laminB2^tail^ expression in “Released” and “Arrested” experimental conditions stained for lamin B1 and GFP and NucBlue to detect DNA. 1G-I. Percent of misshapen nuclei, percent of nuclei with > 10 GFP^+^ foci, and excess nuclear perimeter from GFP-laminB2^tail^-expressing U2OS cells shown in 1F were analyzed as described above. n=3, N=100.

To test this stepwise model, we compared nuclear morphology in dividing and non-dividing cells. Treatment with ribonucleotide reductase inhibitors, hydroxyurea (HU) or thymidine, reversibly arrests cells at S-phase or the G1/S boundary (Wang, 2022), with mitosis peaking ∼12 hours post-release in U2OS cells (Apraiz et al., 2017). Here, U2OS cells were synchronized using hydroxyurea (HU) for 24 hours. Cells were then either maintained in HU to prevent mitosis or released to allow division over the next 24 hours; in both conditions, doxycyline was added for 6-hours to induce GFP-progerin (Fig. 1B). In the absence of doxycycline, fewer than 3% of cells - whether arrested or released - had misshapen nuclei (as measured by a nuclear convexity of <0.945; Fig. 1C). Arrested cells expressing GFP-progerin showed a modest but statistically significant increase in nuclear dysmorphia compared to controls. However, the most pronounced nuclear lobulations were observed in GFP-progerin-expressing cells released from the HU arrest, with over 50% exhibiting misshapen nuclei (Fig. 1C). Conversely, GFP⁺ foci were more prevalent in arrested cells (>60% had at least one focus; 20% had >30 foci) than in released cells (Fig. 1D). We also observed some additional nuclear phenotypes, such as tubules, INM-derived tube-like structures (Arii et al., 2020), and inclusions (Fig. S1B).

Incomplete synchronization may explain the modest increase in lobulated nuclei seen in arrested cells and the presence of foci in released cells, but it is also likely that foci eventually arise de novo in lobulated, post-mitotic nuclei. These phenotypic patterns were also observed in HeLa and hTERT RPE-1 cells (Fig. S1C-D), the latter being a non-transformed line.

GFP-progerin expression and HU-induced cell cycle arrest each increased nuclear area and nuclear perimeter compared to controls (Fig. S1E-F). However, the nuclear perimeter was disproportionately increased in GFP-progerin-expressing under released conditions, as shown by the excess perimeter ratio (Fig. 1E), defined as the ratio of the observed nuclear perimeter to that of a perfect circle with the same area. Released GFP-progerin-expressing cells exhibited an excess perimeter ratio twice that of controls, whereas arrested cells showed no significant change regardless of progerin expression.

### The farnesylated tail domain of laminB2 recapitulates progerin-induced NE phenotypes

Elevated levels of persistently farnesylated, NE-targeted protein such as progerin challenge homeostatic mechanisms that maintain NE architecture. The importance of this dysregulation to aging is underscored by the observation that farnesyltransferase inhibitors ameliorate many HGPS-linked nuclear phenotypes (Bikkul et al., 2018; Foo et al., 2024; Glynn and Glover, 2005; Gordon et al., 2012) and, further, that a non-farnesylatable version of progerin does not cause progeria in a knock-in mouse model (Yang et al., 2011). Different types of NE deformation have been reported to arise when farnesylated NE-targeted proteins are elevated. In some cases, multi-lamellar structures emanating from the INM have been documented (Prüfert et al., 2004; Ralle et al., 2004).

To test whether the two stages of NE phenotype described above (Fig 1B-E) are specific to progerin or represent a broader effect of excess farnesylated lamins, we generated a doxycycline-inducible laminB2^tail^ expression system. Expression of this GFP-tagged, minimal farnesylated domain was sufficient to recapitulate the expanded nuclear perimeter and lobulated nuclear morphology in dividing cells. In arrested cells, laminB2^tail^ accumulated at discrete GFP^+^ NE foci, similar to those seen in progerin, albeit somewhat different in size and frequency, without inducing global contour changes. These findings suggest this two-step NE remodeling pattern is a general principle of adaptation to excess farnesylated proteins at the INM.

### Progerin induces inner nuclear membrane (INM) expansion and invagination

To investigate the nature of the foci formed as an initial consequence of GFP-progerin expression, we examined the localization of nuclear envelope markers, using the LEM-domain protein Emerin (Pawar and Kutay, 2021) to mark the INM and Nesprin2 (Padmakumar et al., 2005), a component of the linker of nucleoskeleton and cytoskeleton (LINC) complex, as an ONM marker. In control arrested cells, Emerin and Nesprin2 were evenly distributed along the nuclear periphery. However, in arrested cells treated with doxycycline to induce GFP-progerin, Emerin accumulated at GFP⁺ foci, while Nesprin2 localization remained unchanged (Fig. 2A). Thus, the GFP^+^ foci appear to be membranous in nature, with selective involvement of the INM.

**Figure 2.**
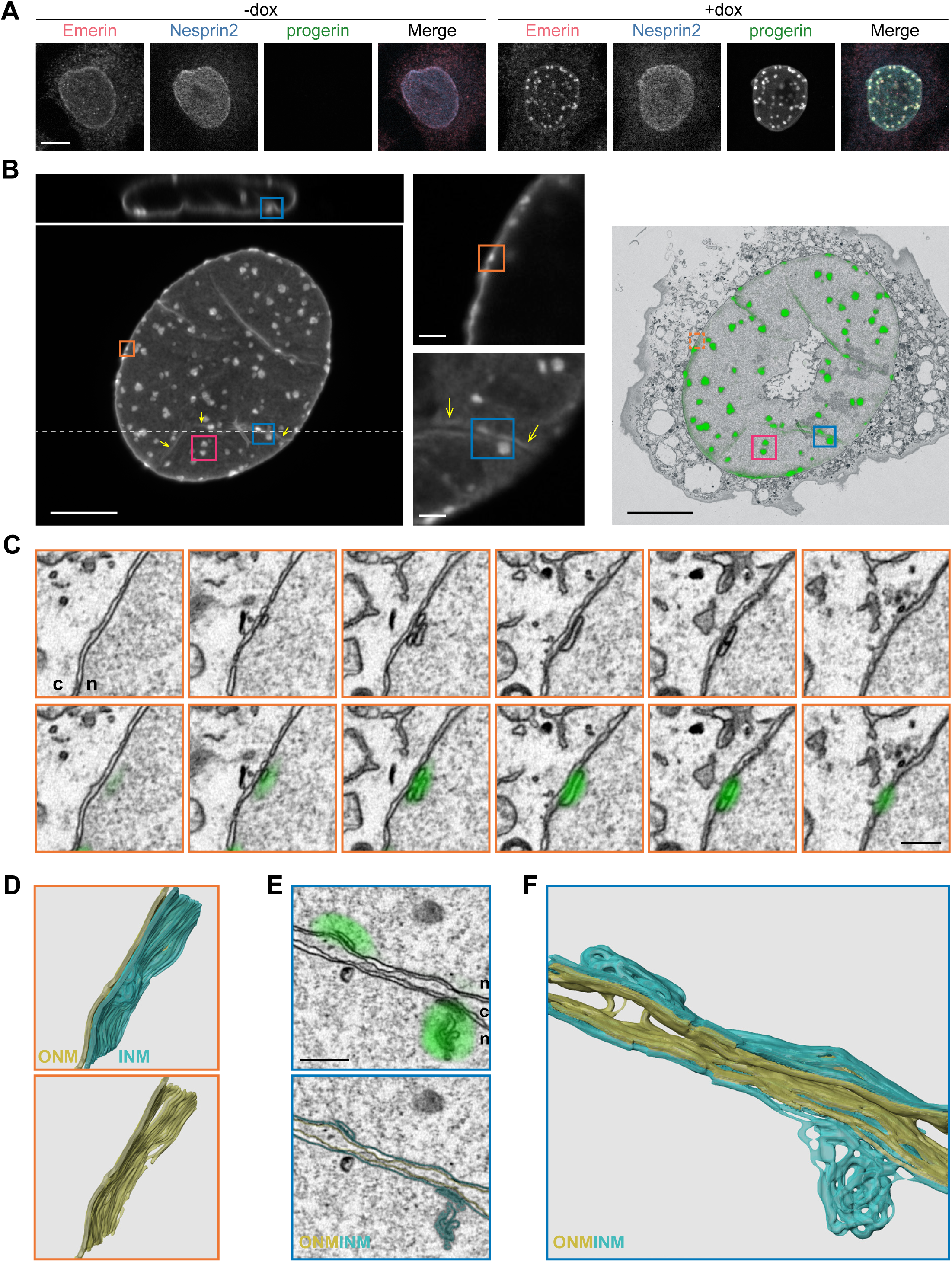
In interphase-arrested cells, progerin forms nuclear envelope-associated foci that correspond to inner nuclear membrane invaginations while the outer nuclear membrane is unaltered. 2A. Confocal images of Emerin, Nesprin2, and GFP-progerin in thymidine-arrested HeLa cells with and without dox-induced GFP-progerin Scale bar = 10 μm 2B. At left, maximum intensity projection of nuclear GFP fluorescence with x-z reconstruction along the dashed line (scale bar = 10 μm) and single confocal z-sections adjacent at right (scale bar = 2 μm). Correlative overlay of 250-nm EM section with GFP-progerin signal (scale bar = 10 μm). Boxes indicate regions of interest annotated in the corresponding electron tomograms (orange and blue, C-F) and Supplemental Movie 1 (magenta); yellow arrows = nuclear fold 2C. Serial EM sections (scale bar = 500 nm) through the correlated region indicated by the orange box in 2B, with and without GFP overlay. n = nucleus; c = cytoplasm. 2D. Tomographic reconstruction of the region annotated by the orange box, rotated to show INM expansion and remodeling at the GFP^+^ focus. ONM = yellow; INM = cyan. 2E. Single EM section (scale bar = 500 nm) through the correlated position indicated by the blue box in 2B, with GFP overlay (top) and nuclear membranes outlined (bottom). On either side of the nuclear fold, multi-layered INM whorls correlate with GFP-progerin foci. ONM = yellow trace; INM = cyan trace; n = nucleus; c = cytoplasm; yellow arrow = nuclear fold. 2F. Tomographic reconstruction of the blue correlated position, rotated to show INM expansion and remodeling at GFP^+^ foci. ONM = yellow; INM = cyan.

To characterize the ultrastructure of these foci, we performed correlative light and electron microscopy (CLEM) and tomography on arrested cells expressing GFP-progerin. Regions of GFP⁺ foci at the lateral and basal nuclear membrane indicated by the orange and magenta boxes, respectively, in Figure 2B correlated with tightly packed, multilayered membranous structures observed by EM (Fig. 2C-D, Movie 1). These membranous structures are derived from the expansion of the INM as there is continuity between the INM and whorls in single EM planes as well as 3D reconstructions (traced in cyan) while the ONM appears largely unaltered (traced in yellow). Invaginations consisting of both inner and outer nuclear membranes have been described (Stiekema et al., 2022) and one such structure, a nuclear fold or cleft, is indicated by the yellow arrows in Figure 2B, F. NE foci in proximity to the nuclear fold within the teal box correspond to sites of GFP-progerin accumulation where INM tubular-lamellar whorls emanate from the fold (Fig. 2E-F). Consistent with this, thin-section electron microscopy of arrested HeLa cells expressing either GFP-progerin or laminB2^tail^ revealed extensive INM invaginations, some forming complex, multi-membranous whorls. The ONM remained structurally intact, and nuclear pore complexes (NPCs) were absent from these invaginations (Fig. S2).

These findings suggest that aberrant accumulation of farnesylated lamins initially drive focal expansion of the INM without significantly altering the ONM. The resulting INM invaginations form localized deformations, thereby maintaining overall nuclear shape and preserving a relatively normal ONM. These NE deformations may correspond to those observed by Goulborne et al. to be dependent on CTP:phosphocholine cytidyltransferase-α (CCTα), which arose with accumulation of farnesylated prelamin A and did so independently of cell division (Goulbourne et al., 2011). We propose that the membrane packing in INM-expanded structures is released upon NE breakdown at mitotic entry. Then, rather than reforming with this topology after division, excess membrane material contributes to the increased nuclear surface area and emergence of the lobulated nuclear morphology characteristic of HGPS.

### A subset of proteins is enriched at progerin-induced INM invaginations

To learn more about the nature of INM expansion that comprises progerin-induced NE foci, we screened a panel of NE-associated proteins for their localization following GFP-progerin induction in arrested cells (Fig. 3A). Immunofluorescence staining revealed that members of the LEM-domain protein family, LEM2, Lap2β, and the above mentioned Emerin, showed strong enrichment at GFP-progerin-induced INM invaginations (Fig. 2A, 3B). In contrast, lamin A and lamin B1 exhibited variable enrichment at these sites (Fig. 3C), consistent with previous reports that type I NR, formed by INM invagination, often lack a complete lamina (Malhas et al., 2011). SUN2, an INM component of the LINC complex, was previously reported to cluster following progerin expression (Vidak et al., 2023) and here was seen to localize robustly to progerin-induced foci. Interestingly, SUN1, also a component of the LINC complex, did not show similar enrichment (Fig. 3D), indicating a level of selectivity in the INM protein repertoire at these sites.

**Figure 3.**
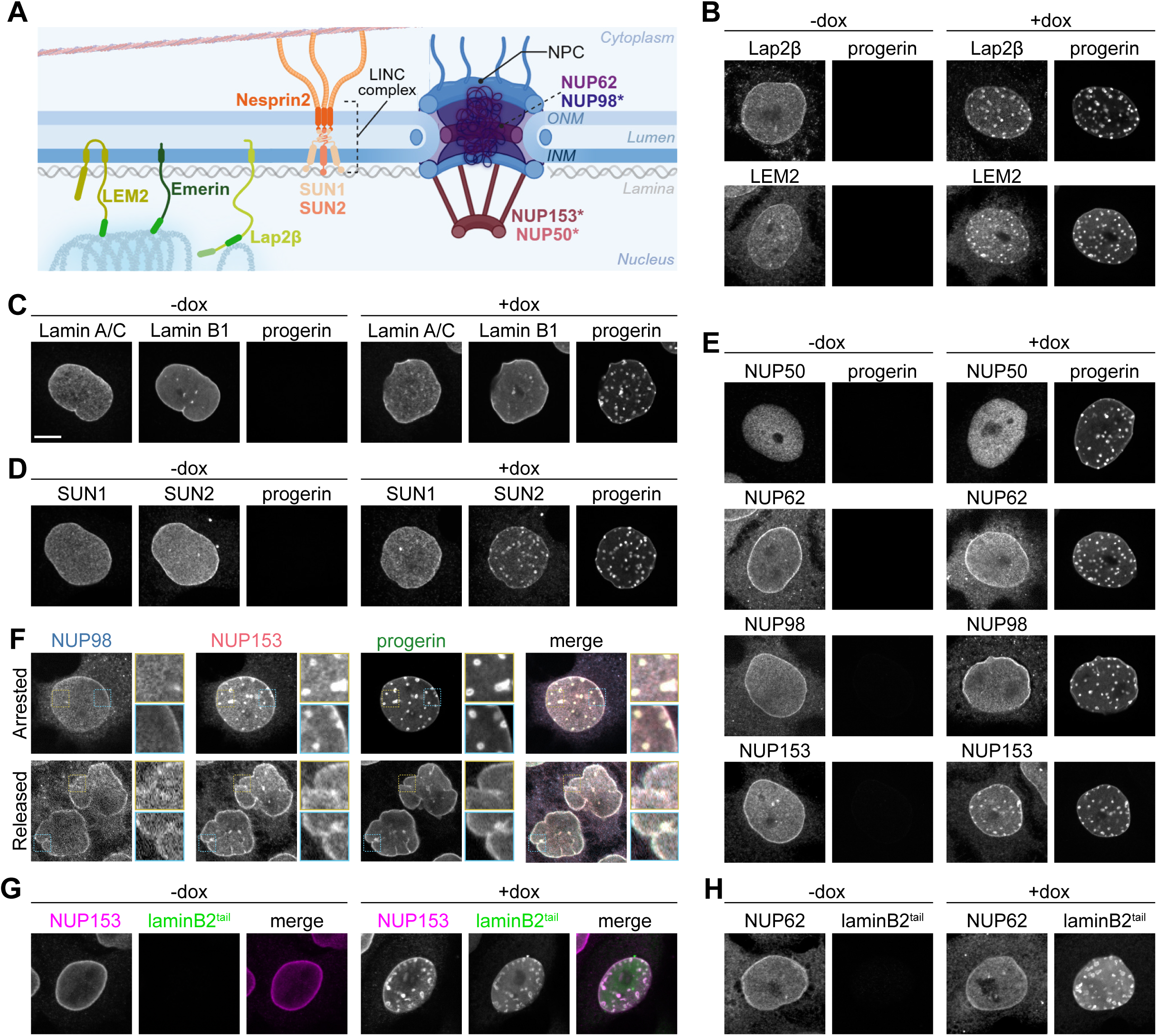
A subset of NE and NPC proteins selectively target to sites of INM invagination. 3A. Schema of NE and associated proteins. 3B-E. Representative confocal images of thymidine- or HU-arrested HeLa cells with dox-inducible GFP-progerin expression, where indicated, showing localization of the annotated NE and associated proteins. Scale bar = 10 μm. 3F. Representative confocal images of HeLa cells expressing GFP-progerin treated under “arrested” or “released” conditions with thymidine treatment (see Figure 1). Colored boxes are 3x blow-ups of regions marked by dotted boxes. 3G-H. Representative confocal images of thymidine-arrested HeLa cells with dox-inducible GFP-laminB2^tail^ expression, where indicated, showing localization of NUP153 and NUP62.

Although nuclear pore complexes (NPCs) were not observed by electron microscopy at progerin-induced INM invaginations (Fig. 2C-F, Fig. S2E-H), we considered whether individual nucleoporins might localize to these regions. Among the nucleoporins tested, NUP153, a dynamic component of the nuclear pore basket (Rabut et al., 2004), was strongly enriched at NE foci, whereas this was not the case for NUP62 nor for NUP98, another mobile nucleoporin that moves between NPCs and the nucleoplasm (Griffis et al., 2002) (Fig. 3E). NUP50, a mobile nucleoporin that is also a constituent of the nuclear pore basket and requires NUP153 for its basket localization (Guan et al., 2000; Hase and Cordes, 2003; Rabut et al., 2004), was also not enriched at these sites. These results raise the possibility that NUP153 accumulates at progerin-induced NE foci in an NPC-independent manner and when we co-stained for NUP153 and NUP98, we found that a population of NUP153 clearly diverges from NUP98 to localize with GFP-progerin (Fig. 3F). While the NR observed by Goulbourne et al. when farnesylated prelamin A accumulated share many attributes with the focal structures observed here in arrested cells (Goulbourne et al., 2011), one difference is their conclusion that NPCs were present. Although the membranous sites of GFP-progerin accumulation detected here may not strictly exclude nuclear pores, we do not think NPCs are prevalent based on our electron microscopy and immunofluorescence results.

We questioned whether selective enrichment of NUP153 could be due to alteration of its overall levels; however, levels of this nucleoporin or others in this panel did not vary with GFP-progerin expression in either arrested or released conditions (Fig. S3A). Of note, the selectivity we observed in arrested cells, where Nesprin2, NUP98, NUP62, and SUN1 did not co-localize with GFP-progerin at NE foci, was not the case when cells were released to divide. Rather, these NE associated proteins co-accumulated at progerin-induced NE folds and lobules, resulting in an uneven localization along the nuclear rim (Fig. 3F; Fig. S3B-E).

To better understand requirements for enrichment at sites of INM expansion (GFP^+^ foci in arrested cells), we probed the localization of representative markers in cells expressing laminB2^tail^ under cell cycle arrest. This resulted in the same pattern of selective enrichment, both for nucleoporins (Fig. 3G-H; Fig. S3F) and INM proteins (Fig. S3. G-H). Thus, although several NE proteins interact with lamin A (Al-Haboubi et al., 2011; Clements et al., 2000; Haque et al., 2006; Liang et al., 2011), their targeting to sites of INM expansion is not dependent on direct interaction with focally accumulated progerin/lamin A. Rather, properties of the local membrane structure induced by the presence of progerin or laminB^tail^ could be key to attracting a distinct set of NE constituents. Amphipathic helices in SUN2 (Lee et al., 2023) and NUP153 (Vollmer et al., 2015) have been implicated in recruiting these proteins to specific membrane contexts, pointing to a potential role for direct affinity in targeting proteins to the tightly packed, multilamellar membrane structures in GFP-progerin-expressing cells.

### NUP153 promotes progerin-induced INM invaginations

To determine whether NUP153 is merely passively attracted to sites of progerin-induced NE deformation or plays a role in their formation and/or stability, we depleted NUP153 in arrested cells expressing GFP-progerin using two independent siRNAs. Based on prior observations that si153-1 is more potent than si153-2 (Mackay et al., 2009), we adjusted their concentrations to 1 nM and 10 nM, respectively, to achieve comparable levels of NUP153 knockdown (Fig. 4A-B). NUP153 depletion led to a significant reduction (averaging 40% for both siRNAs) in the number of GFP^+^ NE foci compared to control siRNA-treated cells (Fig. 4C-D; Fig. S4). Given NUP153’s established role in nucleocytoplasmic transport, including mRNA export and protein import, we considered whether the observed reduction in NE foci could be attributed to impaired import of GFP-progerin or its reduced overall expression. Quantification of nuclear GFP-progerin intensity by immunofluorescence microscopy revealed no significant difference between control and NUP153-depleted cells (Fig. 4E). Furthermore, the nuclear-to-cytoplasmic ratio of GFP-progerin was unaffected by NUP153 knockdown, indicating that its nuclear import was not impaired (Fig. 4F).

**Figure 4.**
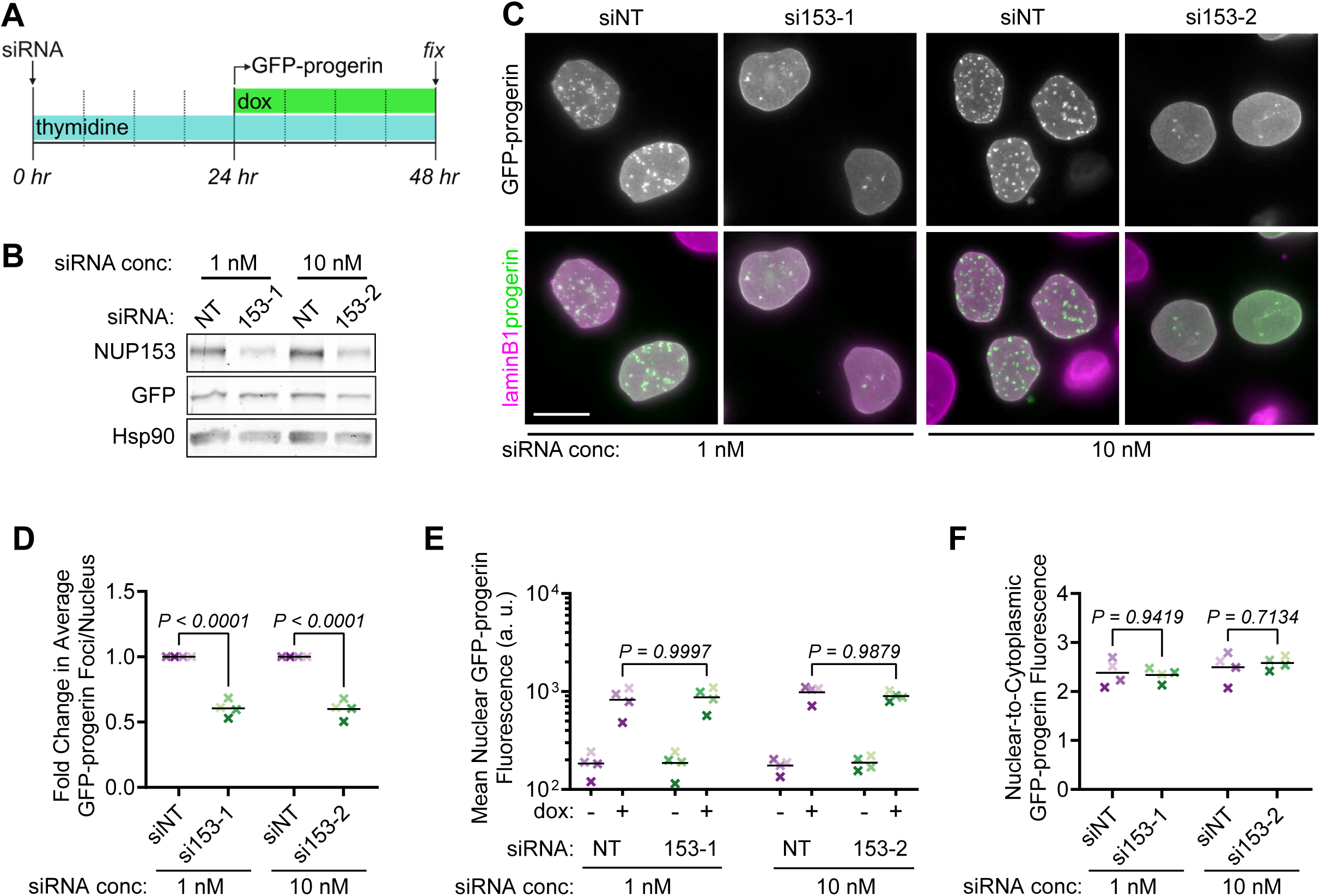
Nup153 depletion attenuates focal INM structures induced by GFP-progerin without disrupting the expression or transport of progerin. 4A. Schema of experimental timeline of HeLa cells treated with siRNA, thymidine, and doxycycline. 4B. Representative western blot analysis of cells treated with 1nM si153-1 and 10nM si153-2 or non-targeting (NT) siRNA. Samples are all +dox and parallel to replicate #4. 4C. Representative widefield images of replicate #4, with lamin B1, tubulin, and GFP detected, as indicated. Scale bar = 20 μm. 4D. Fold change in average GFP^+^ foci per nucleus with Nup153 knockdown compared to its respective control. Data was analyzed using paired one-way ANOVA with post-hoc Tukey’s test. 4E. Mean nuclear GFP-progerin intensity. Data was analyzed using two-way ANOVA with post-hoc Tukey’s test. 4F. Nuclear-to-cytoplasmic ratio of mean GFP-progerin intensity in dox-treated cells. Data was analyzed using paired one-way ANOVA with post-hoc Tukey’s test. Bars represent the mean of four experiments. Each X represents a technical replicate. n=4, N=100.

These findings suggest that NUP153 promotes NE foci formation through a mechanism independent of a canonical role in nucleocytoplasmic transport. A recent study demonstrated that lipid-binding nucleoporins are essential for stabilizing transient nuclear membranes during meiosis (El Mossadeq et al., 2025). Even at interphase, NUP153 depletion was shown to reduce the formation of type II NR associated with DNA-double stranded breaks (DSBs) while depletion of other nucleoporins critical for nucleocytoplasmic transport had no effect on DSB-associated type II NR formation (Shokrollahi et al., 2024). This raises the possibility that NUP153, known to be highly mobile in interphase, has a specialized function in the formation and stabilization of NE invaginations potentially via its amphipathic helix. Consistent with this, NUP153 was previously found to promote tubular-lamellar structures when overexpressed, independent of progerin (Bastos et al., 1996).

Overall, this study demonstrates that nuclear phenotypes arising in the presence of progerin are molded by a combination of mechanisms that initially accommodate the aberrant accumulation of a farnesylated protein at the INM, followed by alterations that occur at the time of subsequent nuclear assembly. Recognition of these distinct stages yields insight into the heterogeneity in cell-based phenotypes following progerin expression and may point to distinct steps to target in designing interventions. The appreciation of stepwise alterations in the NE also provides a new framework for understanding disease etiology and mechanisms that contribute to age-associated changes in nuclear structure-function. More broadly, consideration of the influence of cell division on nuclear dysmorphia may help to parse when and where modulators of nuclear shape (Maheshwari et al., 2021; Schibler et al., 2023) exert their effect.

## MATERIALS AND METHODS

### General cell culture and cell lines

All cell lines used in this study were derived from U2OS (RRID: CVCL_0042), HeLa, or hTERT RPE-1 (RRID: CVCL_4388) cells and confirmed by STR profiling. HeLa cells were kindly provided by Maureen Powers (Emory University), and hTERT RPE-1 cells were provided by Bruce Edgar (University of Utah). Cells were maintained in DMEM supplemented with 10% fetal bovine serum (FBS; Atlas Biologicals) at 37 °C with 5% CO₂. Cultures were routinely tested and confirmed negative for mycoplasma.

Inducible GFP-progerin and GFP-laminB2^tail^ plasmids were constructed using a pLVX-TetOne-GFP backbone containing either a puromycin (PuroR) or G418 (NeoR) resistance cassette. Progerin was generated by Douglas Mackay using TOPO TA cloning. The laminB2^tail^ construct was generated by amplifying the tail domain of lamin B2 (aa399-620). Lamin A and lamin B2 cDNA plasmids were obtained from Bob Goldman (Northwestern University). Constructs were verified by whole plasmid sequencing.

Stable Tet-inducible GFP-tagged cell lines were established by transfecting HeLa and U2OS cells with puromycin-selectable plasmids and hTERT RPE-1 cells with G418-selectable plasmids using Lipofectamine 3000 (Invitrogen) according to the manufacturer’s instructions. Single colonies were isolated and expanded. Doxycycline concentration and treatment duration was optimized for each cell line and experiment (100 ng/mL to 1 µg/mL for 6 h to 24 h, as indicated).

### Cell cycle synchronization

Cells were synchronized at the G1/S boundary by incubation in complete medium containing either thymidine or HU (H8627, Sigma-Aldrich) for 24 h. Arrested cells were treated with media containing thymidine or HU after 24 h to maintain arrest. Released cells were washed thoroughly with PBS and then incubated in fresh medium for 24 h prior to fixation or harvest.

Concentrations were optimized for each cell line: 2 mM thymidine/HU for hTERT RPE-1 and HeLa, and 1 mM for U2OS.

### RNA interference

HeLa cells were transfected with the following siRNA using Lipofectamine RNAiMAX (Invitrogen) and a final concentration of 1 nM or 10 nM siRNA oligonucleotides: si153-1 (Harborth et al., 2001), si153-2: GGACUUGUUAGAUCUAGUU (Mackay et al., 2009). A scrambled version of 153-1, siNT: GCAAAUCUCCGAUCGUAGA (Mackay et al., 2009), was used as a control treatment. Cells were incubated with media containing 2 mM HU and transfection mix for 24 h. Then, the media was replaced with fresh HU-containing medium for an additional 24 h before assessment.

### Live cell imaging

For time-lapse imaging, U2OS cells with a stably integrated dox-inducible GFP-progerin cassette were plated on 35 mm glass-bottom dishes (MatTek, P35G-1.5-20-C) in Leibovitz’s L-15 medium (ThermoFisher) supplemented with 10% FBS. Cells were treated with 500 ng/mL doxycycline and 100 nM SiR-Tubulin with 1 µM Verapamil (Spirochrome) to visualize microtubules. DNA was stained using NucBlue Live ReadyProbes (ThermoFisher). Imaging was performed at 37 °C with 5% CO₂ using a Nikon Eclipse Ti spinning disk confocal microscope equipped with a 60× objective and VisiView software. Cells were imaged every 30 min for 7 h.

To evaluate NE alterations with GFP-progerin expression in arrested cells, U2OS cells were plated as above in imaging medium containing 1mM HU for 24 h, then treated with 100 ng/ml dox for 8 h in HU-containing medium. Cells were then imaged using the Nikon Eclipse Ti spinning disc confocal (60× objective, VisiView) with a 0.2 μm step size between z-slices.

### Immunofluorescence (IF)

Cell were plated on coverslips in 24-well plates and fixed with either −20°C methanol (5–7 min) or 4% paraformaldehyde (PFA) in PBS (15 min, RT). PFA-fixed cells were permeabilized with 0.5% TritonX-100 in PBS for 5 min. Cells were blocked in 3% FBS and 0.05% TritonX-100 in PBS for 40-60 min. Primary and secondary antibody incubations were performed in blocking solution for 1 h at room temperature. The primary antibodies and dilutions used are anti-Emerin

(1:500; 10351-1-AP; Proteintech), anti-GFP (1:1000-1:2000; ab13970; abcam), anti-LaminA/C (1:200; 4777; Cell Signaling), anti-LaminB1 (1:1000; ab16048; abcam), anti-LEM2 (1:1000; 29406-1-AP; Proteintech), anti-Nesprin2 (supernatant; 10H8; a kind gift from Brian Burke), anti-Nup50 (1:500; Bethyl Laboratories), anti-Nup62 (1:200; 610497; Transduction Laboratories), anti-Nup98 (1:200; ab50610; abcam), anti-Nup153 (1:1000; ab96462; abcam), anti-Nup155 (1:500; GTX120945; Genetex), anti-SUN1 (1:250; ab124770; abcam), anti-SUN2 (supernatant; 3.1E; a kind gift from Brian Burke), and anti-tubulin alpha (1:500; MCA77G; Bio-Rad). The secondary antibodies used are AlexaFluor-conjugated antibodies (ThermoFisher). Coverslips were mounted on slides using ProLong Glass Antifade Mountant with NucBlue (Invitrogen).

### IF image acquisition and analysis

Imaging was performed using either spinning disk confocal microscopy (Nikon Eclipse Ti, 60× objective; VisiView software) or widefield microscopy (Zeiss Axioskop 2, 63× objective; Zen Blue software). XZ and YZ cross-sections were reconstructed from confocal image stacks using Fiji/ImageJ. Representative micrographs were processed linearly using Adobe Photoshop (Adobe Systems Inc.) with the adjustment layer/levels tool to adjust contrast and channel intensity. All quantitative analyses were performed on unprocessed images.

Widefield images were analyzed using Zeiss ZenBlue image analysis module. Nuclei were segmented as regions of interest (ROIs) using laminB1 staining. For each ROI, nuclear area, perimeter, and convexity were measured as well as mean pixel intensity in the GFP channel. GFP^+^ nuclei were identified using a threshold determined from untreated controls, and only GFP^+^ nuclei were evaluated under doxycycline-treated conditions.

Nuclear dysmorphia was evaluated using nuclear convexity, an algorithm that calculates the ratio of the convex hull perimeter, formed by connecting the outermost points of the nucleus, to the nuclear perimeter. Nuclei with convexity < 0.945 were classified as misshapen and the percentage of misshapen nuclei was calculated for three independent experiments.

GFP^+^ NE foci were manually annotated by identifying round, dense nuclear puncta with brighter GFP signal than the surrounding NE and automatically counted by the ZenBlue image analysis module for each ROI.

Excess perimeter was used to assess nuclear surface expansion by comparing the perimeter of the nucleus to that of a circle of equal area as previously described (Wang et al., 2024) using the formula:

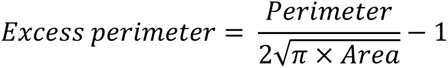

Nuclear-to-cytoplasmic GFP-progerin intensity ratios were calculated using nuclear ROIs and a 12-pixel-wide ring surrounding each nucleus for cytoplasmic measurements. Statistical analyses and graph generation were performed in GraphPad Prism.

### Immunoblotting

Cells were harvested by scraping and lysed in RIPA buffer (150 mM NaCl, 10 mM Tris pH 7.4, 1% deoxycholate, 1% Triton-100, 0.1% SDS) supplemented with 0.5 mM EDTA and HALT protease and phosphatase inhibitor cocktail (ThermoFisher). Samples were normalized using Pierce BCA protein assay kit (ThermoFisher). Samples were resolved by SDS-PAGE and transferred to PVDF membranes by wet transfer. Membranes were blocked in either Intercept (TBS) Blocking Buffer (LICORBio) or 5% non-fat milk in TBS-T and probed with primary antibodies against GFP (1:5000; 632380; Clontech), Hsp90 (1:2000; 13171-1-AP; Proteintech), Vinculin (1:20000; V9131; Sigma), NUP62 (1:1000; GTX102359; GeneTex), NUP98 (1:500; ab179894; abcam), and NUP153 (1:1000; ab96462; abcam), followed by IRDye-conjugated secondary antibodies (LICORBio). Blots were imaged using an Azure Sapphire FL Molecular Scanner (Azure Biosystems) and analyzed with Fiji/ImageJ.

### Transmission electron microscopy

HeLa cells expressing GFP-progerin or laminB2^tail^ were arrested using 2 mM thymidine and treated with 1 µg/mL doxycycline as indicated prior to fixation. Cells were fixed for 10 min at RT in 2.5% glutaraldehyde and 1% paraformaldehyde prepared in 0.1 M sodium cacodylate buffer supplemented with 2.3% sucrose and 8 mM CaCl₂. After fixation, cells were scraped, pelleted in fresh fixative, and post-fixed for 1 h in 2% osmium tetroxide in the same buffer. Samples were rinsed with filter-sterilized deionized water, stained with uranyl acetate, and then dehydrated through a graded ethanol series before being embedded in resin. Ultrathin sections (0.5–1.0 µm) were cut using a Leica EM UC6 ultramicrotome and mounted on copper grids. Sections were sequentially stained with uranyl acetate and Reynold’s lead citrate, then imaged using a JEM-1400Plus transmission electron microscope (JEOL) operating at 120 kV. Electron micrographs were adjusted for brightness in a parallel manner for the figures.

### Correlative light and electron microscopy (CLEM)

U2OS cells stably expressing Tet-inducible GFP–progerin were incubated in FBS-free medium containing LipiBright 650 SMCy5.5 (MG12; Cytoskeleton, Inc.) for 8 h, following the manufacturer’s instructions, in order to use lipid droplets as fiducial markers. After incubation, cells were washed twice with FBS-free medium and re-plated onto 35 mm imaging dishes with gridded coverslips (P35G-1.5-14-C-GRD; MatTek) in complete medium supplemented with 10% FBS for 24 h. Cells were then arrested with 1 mM hydroxyurea (HU) for 24 h. Subsequently, the medium was replaced with fresh medium containing 1 mM HU and 100 ng/mL doxycycline for 8 h. Cells were treated with 60 μM cumate (QM150A-1; System Biosciences) before and after HU addition to induce a LEM2 construct which ultimately was not tracked. Cells were fixed with freshly prepared 2% paraformaldehyde (PFA) in PBS (pH 7.4) for 15 min at room temperature (RT). Light Microscopy (LM) of direct fluorescence (20× and 60× objectives) and brightfield (20× objective) was performed using a spinning disk confocal microscope (Nikon Eclipse Ti; VisiView). Z-stacks were collected at 0.25 μm intervals, and grid patterns were tracked from brightfield images for CLEM alignment.

Following imaging, cells were incubated with freshly prepared fixative containing 3% PFA and 3% glutaraldehyde in PBS (pH 7.4) for 30 min at RT and stored at 4 °C overnight. Samples were then stained and embedded in resin as described (Ewald et al., 2012; Mitchell et al., 2024).

3D spin-milling was performed on a Thermo Fisher Helios Hydra Plasma-focused ion beam (PFIB) system, which operates with an oxygen plasma ion source. A ∼1.5 mm-thick resin slab containing cultured cells on its top surface was mounted onto an SEM stub using silver paste, without sputter coating. The sample was positioned at the system’s eucentric height and the stage was tilted to –34°, resulting in a 4° glancing incidence angle for the FIB. Initially, severe charging of the resin caused the secondary electron detector to saturate and obscure cellular contrast. To mitigate this, a 12 keV, 64 nA oxygen beam was applied broadly across the sample. After ∼5 min of exposure, the charge dissipated, allowing clear visualization of the embedded cells and easy identification of the cell previously imaged by confocal microscopy, aided by the etched grid number transferred from the coverslip.

The spin-milling workflow proceeded as follows. First, the oxygen FIB beam (12 keV, 64 nA) was scanned in a 500 × 40 µm box over the region of interest for 30 seconds to expose a fresh surface. The stage was then compucentrically rotated by 72°, and the milling step was repeated. This sequence was performed five times to complete a 360° rotation, delivering ion flux from multiple azimuthal angles to reduce milling-induced texture. One full rotation constituted a single “z-slice.” Next, the stage was returned to 0° tilt for SEM imaging of the target cell using the retractable backscattered electron detector. These milling and imaging steps were automated using Auto Slice & View (ThermoFisher Scientific) and repeated until the full imaging volume was acquired.

The milling thickness was maintained at ∼40 nm per slice, with an in-plane pixel size of 10 nm. A total of 134 slices were collected to span the entire single-cell volume. Resulting 3D datasets were aligned, reconstructed, and visualized in Amira (ThermoFisher Scientific), which was also used to correlate the EM and LM datasets.

## ACKNOWLEDGEMENTS

We thank Chris Jensen (University of Utah) and Nancy Chandler (EM Core, University of Utah) for technical assistance. We also thank Drs. Natasha Saik (University of Utah) and Natalie Kirkland (Altos Labs) for their helpful suggestions.

