## Supplementary Figures for "Progerin-Induced Nuclear Envelope Remodeling is Shaped by Cell Division and NUP153"

#### SUPPLEMENTARY INFORMATION

#### SUPPLEMENTARY FIGURE LEGENDS

**Supplemental Figure 1. Accumulation of persistently farnesylated lamin constructs, progerin and laminB2$^{tail}$, results in cell cycle stage-specific NE deformation**

A. The addition of a farnesyl group to the C-terminal lamin tail domain anchors these proteins to the inner nuclear membrane. Prelamin A is processed to mature lamin A through proteolytic cleavage of its farnesylated tail. In HGPS, a point mutation promotes the use of a cryptic splice site resulting in an isoform called progerin, that is missing 50 amino acids (aa607-656) containing the cleavage site, resulting in the accumulation of permanently farnesylated progerin. Lamin B2 is permanently farnesylated at its C-terminus. The farnesylated residues of the Lamin B2 tail domain (aa399-620), like Lamin B2, are not subject to proteolytic removal. IF Rod = intermediate filament rod domain; NLS = nuclear localization signal; LTD = lamin tail domain.

B. Schema and representative images of HU-arrested U2OS cells with a stably integrated doxycycline (dox)-inducible GFP-progerin cassette. Cells were incubated with 100ng/ml dox for 8 hours before imaging live using a spinning disc confocal microscope. Z-sections acquired with 0.2 µm step size were used to reconstruct x-z and y-z cross-sectional profiles (boxes) corresponding to the plane at the matching dotted lines; top, x-z cross-sectional profile; right, y-z cross-sectional profile. In addition to NE foci on both apical and basal nuclear membranes (orange box; orange arrows indicate a focus on the basal nuclear membrane), GFP-progerin accumulated at NE-associated spherical inclusions (yellow arrows) as well as tube-like structures (magenta arrows, tube; magenta arrowhead, tube opening) and tubules (cyan arrow) which traverse the length of the nucleus. Step size between representative images = 0.8 µm. Scale bar = 10µm.

C-D. Representative images from widefield microscopy of HeLa (C) and hTERT RPE-1 (D) cells treated with 2 mM HU for 24 hours before treating with 1 µg/ml dox either with (Arrested) or without (Released) HU for another 24 hours. Antibodies were used to detect lamin B1, tubulin, and GFP (progerin) by indirect immunofluorescence and NucBlue used to detect DNA. Scale bar = 20µm

E. Nuclear area in U2OS cells expressing GFP-progerin from experiments described in Figure 1B. Measurements were made from widefield images and only GFP$^{+}$ nuclei were evaluated in dox-treated conditions. Each dot represents a single nucleus, with different shades representing replicates. Each X represents the mean of a technical replicate. n=3, N=100. Data was analyzed using a Two-Way ANOVA with post-hoc Tukey's test.

F. Nuclear perimeter in U2OS cells expressing GFP-progerin from experiments described in Figure 1B. Measurements were made from widefield IF images and only GFP$^{+}$ nuclei were evaluated in dox-treated conditions. Each dot represents a single nucleus, with different shades representing replicates. Each X represents the mean of a technical replicate. n=3, N=100. Data was analyzed using a Two-Way ANOVA with post-hoc Tukey's test.

**Supplemental Figure 2. Progerin and laminB2$^{tail}$ expressing HeLa cells accumulate INM deformations**

A. Representative images of direct GFP fluorescence from parallel widefield microscopy of HeLa cells stably expressing the indicated dox-inducible cassette. Cells were arrested with thymidine and treated with 1 μg/ml dox for 24 hours, where indicated. Scale bar = 20 μm

B-D. Thin section electron micrographs of whole nuclei of untreated control (B), progerin-expressing (C), and laminB2$^{tail}$-expressing (D) HeLa cells reveal regularly-shaped nuclei with smooth ONM. The INM was relatively unperturbed in control cells while cells expressing farnesylated lamins developed INM-associated structures. Scale bars = 5 μm

E-H. Representative electron micrographs of INM-associated structures from cells expressing progerin (teal frame) and laminB2$^{tail}$ (magenta frame). Narrow tubular invaginations of the INM projecting a straight or convoluted path into the nucleoplasm (E, scale bars = 500nm), single-layered membranous whorls (F, scale bar = 500 nm), and multi-layered membranous whorls (G, scale bars = 1 μm; H, scale bar = 500 nm) were observed with progerin and laminB2$^{tail}$ expression but not in untreated cells.

I. The membranous deformations shown in E-H were quantified from thin section electron micrographs of whole nuclei. INM-associated invaginations and whorls accumulated in cells expressing farnesylated lamins but were absent in control cells.

**Supplemental Figure 3. Selective targeting of NE constituent to sites of both focal GFP-progerin and laminB2$^{tail}$ accumulation and proteins that do not enrich at these sites localize alongside GFP-progerin/laminB2$^{tail}$ at nuclear folds and lobulations in nuclei after cell division**

A. Western blot showing levels of GFP-progerin, lamin A, and lamin C as well as NUP62, NUP98, and NUP153 under released and arrested conditions. NUP62, NUP98, and NUP153 expression appears unchanged with progerin expression.

B-D. Representative confocal images of GFP-progerin-expressing HeLa cells under released conditions treated with 1 µg/ml doxycycline for 24 hours. Progerin was detected using a GFP antibody. Most progerin-expressing cells exhibited lobulated nuclei with multiple folds, while a few regularly shaped nuclei contained foci, likely representing cells that had not divided after progerin accumulation. The indicated NE and NPC associated proteins, detected with specific antibodies, were consistently enriched at NE folds but showed selective enrichment at foci. Scale bars = 20 µm.

E. Representative confocal images of GFP-progerin-expressing HeLa cells under released conditions. In untreated control cells, NUP62 colocalized with the INM protein lamin B receptor (LBR) at the nuclear rim. In dox-treated cells, both proteins were enriched at NE folds. Notably, LBR also localized to foci in unlobulated nuclei (yellow boxes, 2x blow-ups), whereas NUP62 did not. Scale bar = 20 µm.

F. Representative confocal images of GFP-laminB2$^{tail}$-expressing HeLa cells under released conditions stained for NUP153. LaminB2$^{tail}$**,** detected using a GFP antibody, localized to both the nucleoplasm and nuclear rim. NUP153 was strongly enriched at the nuclear rim in control cells and at laminB2$^{tail}$-induced NE foci and folds, indicating that NUP153 localization to NE invaginations does not depend on its interaction with a specific lamin. Scale bar = 20 µm.

H-I. Representative widefield images of thymidine-arrested HeLa cells treated with 1 µg/mL doxycycline for 24 h. Similar to progerin-induced NE foci, SUN1 and Nesprin-2 were not enriched at laminB2$^{tail}$-induced NE foci foci, whereas Emerin and SUN2 were.

**Supplemental Figure 4. Focal INM structures induced by GFP-progerin are reduced with NUP153 depletion**

The average number of GFP-progerin foci per nucleus in control and Nup153-depleted cells. Data corresponds to that summarized in 4D. Dots represent single nuclei, with different shades representing replicates; each X represents the mean of a technical replicate. n=4, N=100. Data was analyzed using paired one-way ANOVA with post-hoc Tukey's test.

**Supplemental Figure 5. Full Western blots from indicated figures**

**MOVIE 1**

**GFP-progerin accumulates at discrete sites at the NE resulting in INM-derived structures (related to Fig. 2B)**

CLEM tomogram reconstruction (y-z) of the indicated ROI (magenta box, Fig. 2B) showing the apical and basal nuclear membranes. Two $GFP^{+}$ foci located along the basal nuclear membrane correspond to sites of well-demarcated INM expansion and folding resulting in multi-layered membranous whorls. Movie shown at 25 frames per second. Scale bar = 1 μm.

### Turkmen et al. Supplemental Figure 1

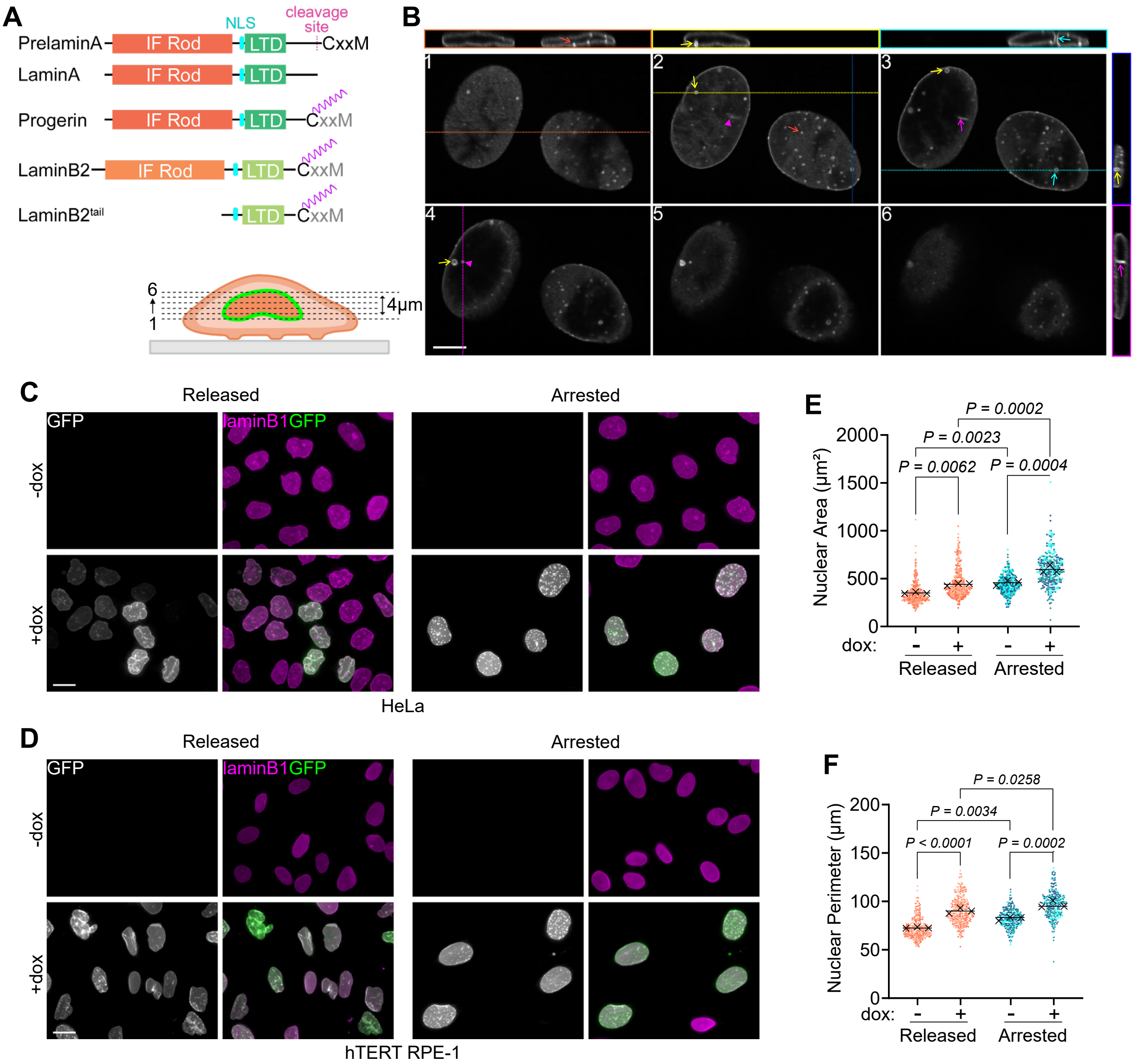

**Turkmen et al. Supplemental Figure 2**

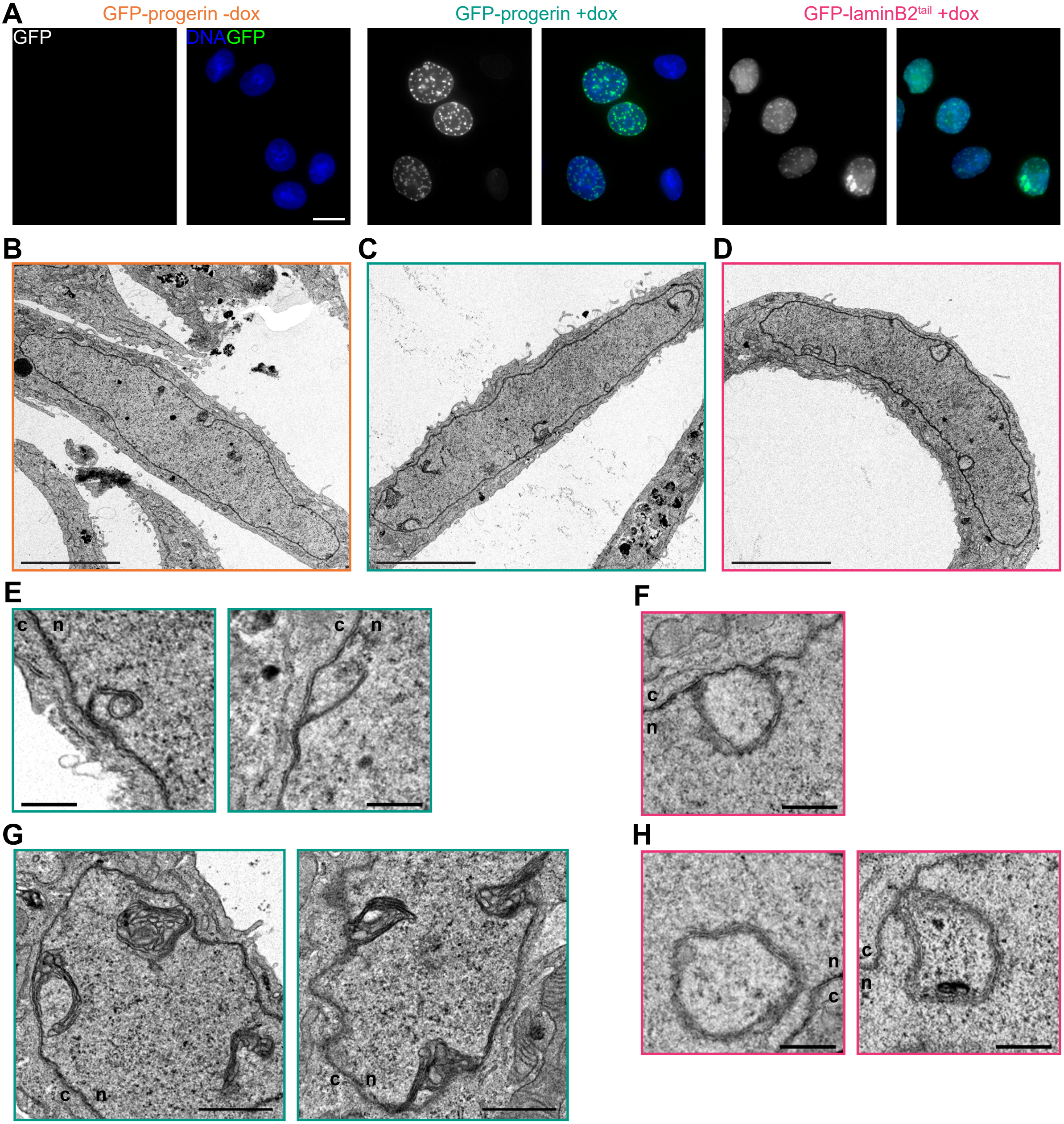


I

| construct | GFP-progerin | GFP-progerin | GFP-laminB2tail |
|---|---|---|---|
| treatment | -dox | +dox | +dox |
| number of cells | 10 | 21 | 20 |
| single-layered INM inv. | 0 | 5 | 2 |
| single-layered INM whorls | 0 | 6 | 27 |
| multi-layered INM whorls | 0 | 24 | 4 |

### Turkmen et al. Supplemental Figure 3

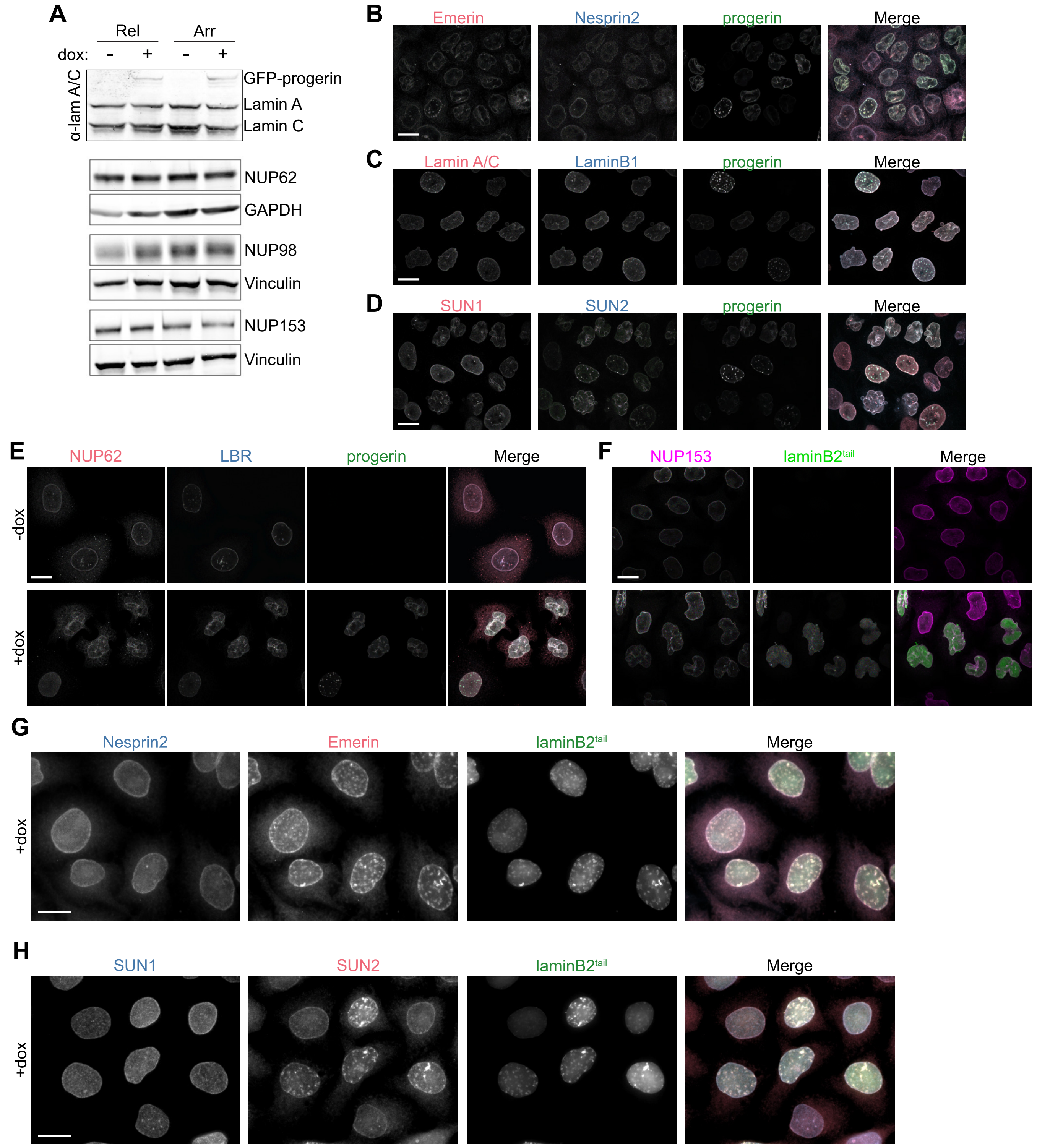

### Turkmen et al. Supplemental Figure 4

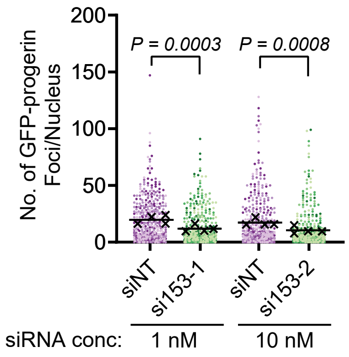

### Turkmen et al. Supplementary Figure 5

#### Figure 4:

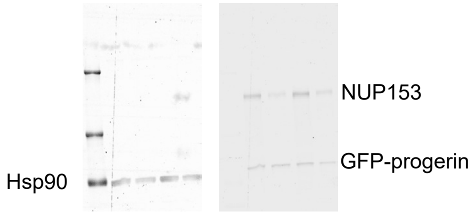


#### Supplementary Figure 3:

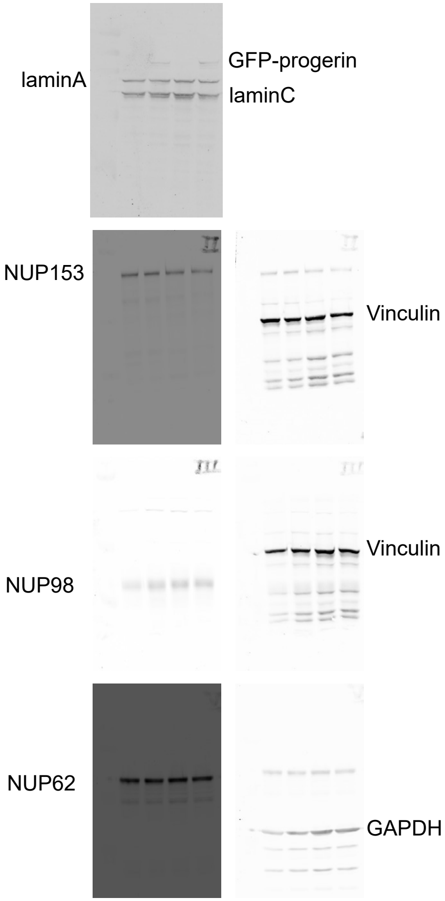
